# Newly identified relatives of botulinum neurotoxins shed light on their molecular evolution

**DOI:** 10.1101/220806

**Authors:** MJ Mansfield, TG Wentz, S Zhang, EJ Lee, M Dong, SK Sharma, AC Doxey

**Author notes:** Equal contribution (MJ Mansfield, TG Wentz). Correspondence should be addressed to A.C.D. and S.K.S.

## Abstract

The evolution of bacterial toxins is a central question to understanding the origins of human pathogens and infectious disease. Through genomic data mining, we traced the evolution of the deadliest known toxin family, clostridial neurotoxins, comprised of tetanus and botulinum neurotoxins (BoNT). We identified numerous uncharacterized lineages of BoNT-related genes in environmental species outside of *Clostridium*, revealing insights into their molecular ancestry. Phylogenetic analysis pinpointed a sister lineage of BoNT-like toxins in the gram-negative organism, *Chryseobacterium piperi*, that exhibit distant homology at the sequence level but preserve overall domain architecture. Resequencing and assembly of the *C. piperi* genome confirmed the presence of BoNT-like proteins encoded within two toxin-rich gene clusters. A *C. piperi* BoNT-like protein was validated as a novel toxin that induced necrotic cell death in human kidney cells. Mutagenesis of the putative active site abolished toxicity and indicated a zinc metalloprotease-dependent mechanism. The *C. piperi* toxin did not cleave common SNARE substrates of BoNTs, indicating that BoNTs have diverged from related families in substrate specificity. The new lineages of BoNT-like toxins identified by computational methods represent evolutionary missing links, and suggest an origin of clostridial neurotoxins from ancestral toxins present in environmental bacteria.

**Significance statement:** The origins of bacterial toxins that cause human disease is a key question in our understanding of pathogen evolution. To explore this question, we searched genomes for evolutionary relatives of the deadliest biological toxins known to science, botulinum neurotoxins. Genomic and phylogenetic analysis revealed a group of toxins in the *Chryseobacterium piperi* genome that are a sister lineage to botulinum toxins. Genome sequencing of this organism confirmed the presence of toxin-rich gene clusters, and a predicted *C. piperi* toxin was shown to induce necrotic cell death in human cells. These newly predicted toxins are missing links in our understanding of botulinum neurotoxin evolution, revealing its origins from an ancestral family of toxins that may be widespread in the environment.

## Introduction

Despite a wealth of detailed knowledge on the structure-function relationship of many bacterial toxins (1–4), the evolutionary origins of important toxins responsible for human disease remain unclear (5, 6). Although bacterial toxins are typically studied in the context of their pathogenesis toward human cells, modern toxins are likely descendants of ancestral proteins that targeted difference substrates, cell types and hosts. By identifying the evolutionary relatives of bacterial toxins in diverse environments (5), it may be possible to not only reconstruct these toxin evolutionary histories, but also identify sequence changes responsible for their unique activities (5–8).

Clostridial neurotoxins, including botulinum neurotoxins (BoNTs) and tetanus neurotoxin (TeNT), the causative agents of botulism and tetanus, are the most deadly toxins known to science with LD_50_ values ranging from 0.1-1.0 ng per kg (9). Clostridial neurotoxins represent serious threats to public health and require constant monitoring by health agencies. Owing to their extreme toxicity, BoNTs are potential bioterrorism agents, and yet also have enormous utility as protein therapeutics (10, 11). BoNTs are produced by *Clostridium botulinum*, a polyphyletic taxon classified solely by the presence of the neurotoxin, and several other *Clostridium* species. Neurotoxin genes reside in distinct gene clusters encoded on the chromosome, plasmids or phages.

The extreme toxicity of BoNT is ultimately due to its structure and function. BoNTs are initially produced as a single polypeptide chain, which is then cleaved by a bacterial protease to result in a light-chain (LC) and heavy-chain (HC) which remain linked by a disulfide bond. A C-terminal receptor-binding domain in the heavy chain (H_CC_), which adopts a ricin-type beta-trefoil structure, targets BoNTs to different neuronal receptors in the host, including SV2 for BoNT serotypes A/D/E/F, and synaptotagmin I/II for BoNT serotypes B/G with polysialogangliosides as co-receptors (12–23). After neuronal binding, BoNTs are internalized within endocytic vesicles. At low pH, the HC translocation domain, which forms an all alpha-helical bundle structure, transports the partially unfolded LC into the cytosol. The BoNT-LC, an ~400 residue N-terminal zinc metalloprotease domain, then cleaves intracellular SNARE proteins including VAMPs, SNAP25, and syntaxin 1 (24–27) to prevent exocytosis of synaptic vesicles, leading to flaccid paralysis (3).

The sophistication of BoNT’s mechanism of action (see Rossetto et al. (3) for a thorough review) raises questions about its evolutionary origins. Recent work describing a divergent homolog of BoNTs in the genome of *Weissella oryzae*, suggests that BoNT-related proteins may not be limited to the *Clostridium* genus (8, 28). As the *Weissella* toxin is the only identified homolog outside of the BoNT family to date, however, it is unclear whether it represents a highly divergent pseudogene, or if it indicates the existence of a larger *bont*-like gene family present in a broader range of bacterial taxa.

Here we present a large-scale bioinformatic screen for putative BoNT-related toxin genes in all available genomes. Our search identified 161 BoNT-related genes in 30 different species. Predicted genes include new candidate toxins from *Chryseobacterium*, *Mycobacterium*, *Streptomyces*, and 19 species of entomopathogenic fungi, all containing signatures of toxin functionality. We identified toxin candidates in the *Chryseobacterium piperi* genome (29), which we resequenced and verified to encode two toxin-containing gene clusters with tandem duplications reminiscent of BoNT-NTNH. Experimental validation of *C. piperi* toxins revealed a candidate toxin with metallopeptidase activity that arrested cell proliferation in human kidney cells. Phylogenetic analysis suggests that the *C. piperi* and other identified toxins are missing links that shed light on the evolution of BoNTs from ancestral protein toxins present in environmental bacteria.

## Results

### Genomic data mining predicts BoNT-like toxins outside of Clostridium

We screened the NCBI Genbank database (March 26, 2017) comprised of 94,396 prokaryotic, 4,123 eukaryotic, and 7,178 viral genomes, for potential homologs of BoNTs. Using PSI-BLAST with selected BoNT sequence queries (see Methods), we detected a total of 309 protein sequences displaying at least partial homology to BoNTs with an *E*-value below 0.001 (Table S1). The dataset includes all known BoNT serotypes, and an additional 161 BoNT-related sequences, all of which are experimentally uncharacterized to date. We performed all-by-all pairwise alignments and clustered the toxins using principal coordinates analysis (PCoA), which revealed three main sequence clusters (Fig. 1*A*). Cluster I includes a large family of putative ADP-ribosyltransferase toxins from entomopathogenic fungi, as well as diphtheria toxin, all possessing partial similarity only to the BoNT translocation domain (17.3% max sequence identity, PSI-BLAST *E*-value = 7 x 10^-^40). Cluster II includes type III effector proteases such as *Escherichia coli* NleD (peptidase family M91) which cleaves host JNK and p38 (30), and which possess remote detectable homology to the BoNT-LC with 14.9% max sequence identity and a PSI-BLAST *E*-value of 4 x 10^-^5 (see Methods).

**Figure 1.**
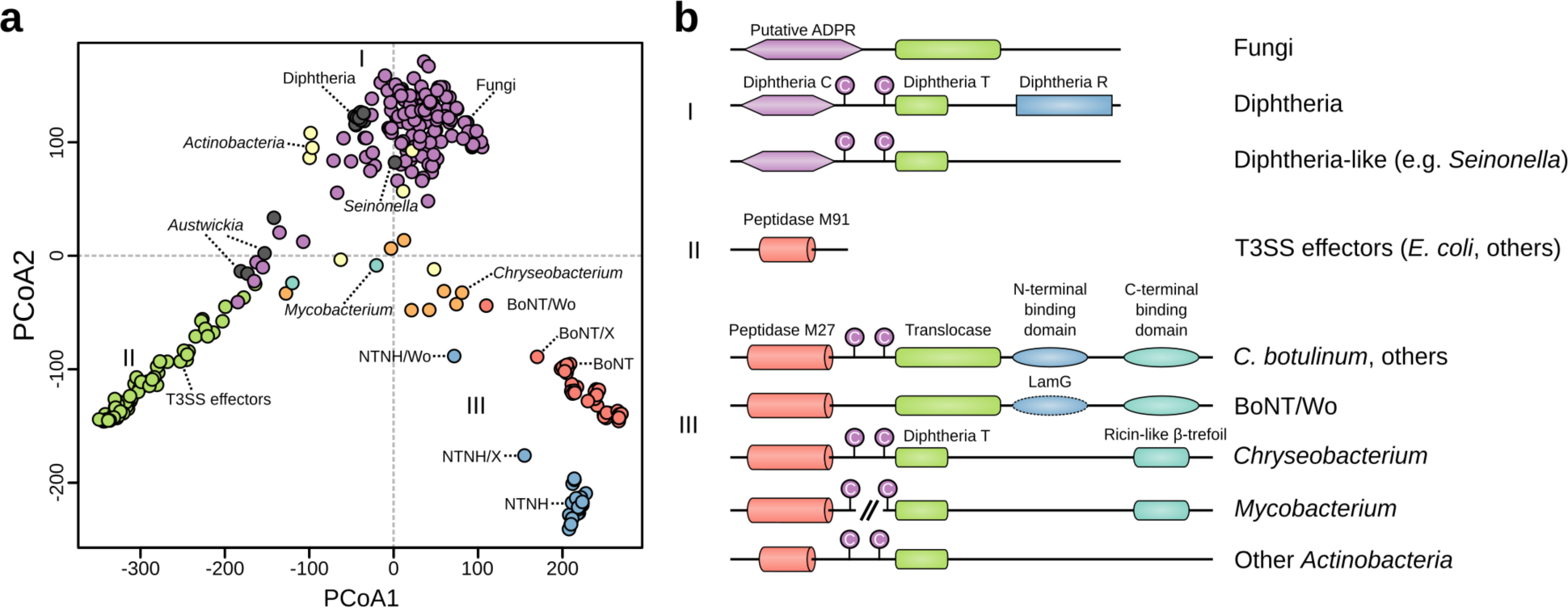
Visualization of sequence and domain architectural similarities between BoNTs and their evolutionary relatives detected by PSI-BLAST. *A*) PCoA ordination of pairwise percent similarities reveals relationships between the sequence families surveyed in this study. Predicted toxins from entomopathogenic fungi cluster distinctly from BoNTs, and associate more closely with diphtheria toxins (cluster I). Type III secretion system (T3SS) effectors group separately (cluster II). BoNT and NTNH form distinct groups, with more divergent relatives such as the *Weissella* “BoNT/Wo” clustering outside them. The next closest relatives are the BoNT-like toxins found in *Chryseobacterium*, followed by putative toxins from other *Actinobacteria* (cluster III), which span the distance between groups, having similarities to each. *B*) Predicted domain architectures of the sequences identified in this survey. Homologs detected by PSI-BLAST include proteins with BoNT-like domain architectures (e.g., *Weissella* “BoNT/Wo”) or share only a single domain (e.g., the fungal family). For the sake of simplicity, only a single sequence was chosen to represent the subgroups, and domain structures can vary further within the groups. Notably, genes encoding *M. chelonae* BoNT-like proteins appear to be split into two components (one encoding the “light chain” M27-like peptidase, and the second encoding the “heavy chain” translocation domain and receptor-binding domain).

The lower right cluster (III) contains BoNTs, NTNHs, the *Weissella* toxin and several uncharacterized proteins (Fig. 1*A*) that share multiple domains in common with BoNTs (Fig. 1*B*, Fig. S2) and are therefore of considerable interest (8, 31). Among the uncharacterized proteins are nine partial and full-length homologs from the genome of *Chryseobacterium piperi*, two from *Mycobacterium chelonae*, and five from other *Actinobacteria*. Some of these organisms are associated with disease; *Chryseobacterium* species are opportunistic pathogens (32, 33), *Acaricomes phytoseiuli* is a pathogen of mites (34), and *Mycobacterium chelonae* is a human pathogen associated with skin, soft tissue, and bone infections (35). We termed these proteins “BoNT-like toxins” given their evolutionary distance to BoNTs (Fig. 1*A*) and similarity of domain architecture (Fig. 1*B*). As shown for a representative protein from this group (putative *Chryseobacterium* toxin, “Cp1”, NCBI accession number WP_034687872.1), these proteins possess a BoNT-like three domain architecture composed of a predicted metalloprotease domain, central translocation domain and C-terminal ricin-type beta-trefoil domain (Fig. 1*B*), each of which are analyzed further in detail below. Despite possessing detectable homology spanning multiple domains (Fig. S2), BoNT-like toxins have low sequence identity to BoNTs (17% identity between Cp1 and BoNT/A compared to >=28% identity between BoNT family members) indicative of a distant evolutionary relationship.

### Chryseobacterium toxins are a sister lineage to BoNTs

Putative protease domains from BoNT-like toxins were aligned to the BoNT-LC to perform sequence, structural, and phylogenetic analysis (Fig. 2, Fig. S3, S4). Phylogenetically, BoNTs grouped into a distinct clade with BoNT/X and *Weissella* toxin forming divergent early branching lineages (Fig. 2*A*, Fig. S4). A sister lineage consisting of the four *C. piperi* homologs and one *M. chelonae* homolog next grouped with the BoNT family with high statistical confidence (98% boostrap support) (Fig. 2*A*, Fig. S4). Additional toxins from *Streptomyces* grouped next with this cluster, and finally a large clade of distantly related, type III effector proteases (M91 family) (Fig. 2*A*). Despite some variable segments and low sequence identity (BoNTA1/CP1: 17.9%), the protease domains from *C. piperi* and other BoNT-like toxins possess detectable homology to the BoNT-LC (bl2seq *E*-value = 2 x 10^-^6 between Cp1 and BoNT/A1) and conserve key functional residues found in BoNTs (Fig. 2*B*). These residues include: the critical HExxH zinc-coordinating active site motif; the third zinc ligand Glu-261; the functionally important Glu-350 which shapes active site fine structure, the active site stabilizing motif R363-x-x-Tyr366 (36), and the cysteine residues that form the disulfide bridge between BoNT LC and HC (37) (Fig. 2*B*).

**Figure 2.**
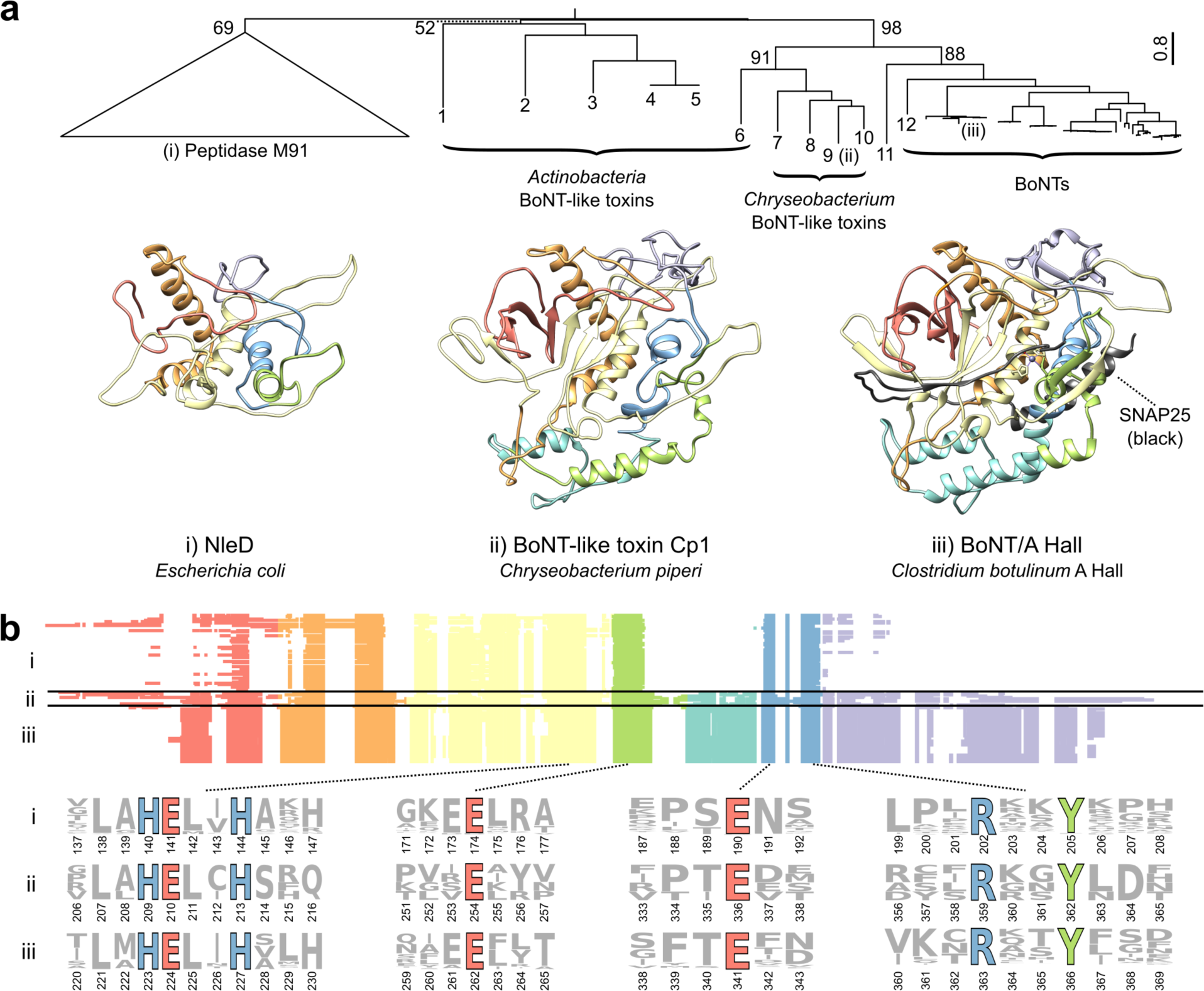
Comparison of the BoNT-LC with homologous domains from BoNT-like toxins and M91 family type III secretion system effectors. *A*) Phylogenetically, BoNTs form a statistically supported clade with BoNT-like toxins distinct from peptidase M91. Structural comparison of BoNT/A (PDB 3BTA) (iii) with structural models of *Chryseobacterium* Cp1 (ii) and *E. coli* NleD (i) reveals two regions that are unique to BoNTs and BoNT-like proteins: the lower alpha-helical region, which interacts directly with SNARE substrates, and the C-terminal region that plays a role in catalytic product removal. *B*) The protease domains of type III secretion system effector proteases (i), BoNT-like toxins (ii), and BoNT-LCs (iii) have key conserved sequence features. These include the HExxH zinc-coordinating and catalytic residues, the third zinc ligand E261, and the active site-refining E350 and RxxY motif. As depicted in the multiple alignment, two insertion regions appear unique to BoNTs and BoNT-like proteins, shown in teal and purple respectively. The identities of proteins labeled 1-12 are: 1 - WP_055473237.1, *Streptomyces pathocidini*; 2 - GAO13068.1, *Streptomyces* sp. NBRC_110027; 3 - WP_083906476.1, *Acaricomes phytoseiuli*, 4 - WP_037712107.1, *Streptomyces* sp. AA4; 5 - EFL04418.1, *Streptomyces* sp. AA4; 6 - WP_070931163.1, *Mycobacterium chelonae*; 7 - WP_034681281.1, *C. piperi*; 8 - WP_034687877.1, *C.* piperi; 9 - WP_034687872.1, *C. piperi*; 10 - WP_034687193.1, *C. piperi*; 11 - WP_027699549.1, *Weissella oryzae* SG25; 12 - BAQ12790.1, *Clostridium botulinum* str 111 (BoNT/X).

Consistent with phylogenetic analysis, the predicted structure of *C. piperi* toxins are most similar to BoNT-LC (7.0Å RMSD versus 12.0Å for *E. coli* NleD) (Fig. 2*A*). One insertion is common to BoNT-LCs and BoNT-like toxins, and absent in NleD and other M91 proteases (Fig. 2*A,B*), which makes extensive contacts with SNAP25 (51 inter-residue contacts < 2 Å) and VAMP2 (91 inter-residue contacts < 2Å) in BoNT co-crystal complexes (Fig. S5). This insertion may therefore have contributed to an ancestral shift in substrate specificity between M27 and M91 protease families. A second C-terminal insertion common to BoNT-LC and BoNT-like toxins (Fig. 2*A,B*) forms part of the hydrophobic SNAP25 binding pocket (38), and has been shown to mediate catalytic activity and product removal (39).

In addition to LC conservation, *Chryseobacterium* and other BoNT-like toxins possess significant similarity to the BoNT translocation domain (22% identity, bl2seq *E*-value < 1 x 10-^5^), particularly across the region 593-686 in BoNT/A1 (Fig. S6), which forms a critical channel-forming amphipathic alpha helical motif (40–42). This region common to BoNTs and BoNT-like toxins is also highly similar to the membrane-inserting residues 286-325 of diphtheria toxin (helices TH5-TH6/TH7, PDB ID 4AE0) (43), consistent with a structural prediction for this region made by PHYRE (Fig. S2). In particular, the motif [K/R]x(8)PxxG is universally conserved among the translocation regions of all the identified toxins in our screen except the single-domain M91 family proteases (Fig. 1*B*, Fig. S6), highlighting it as a key sequence signature of translocation function.

Lastly, following the translocation domain, BoNT-like toxins possess a receptor-binding domain that is predicted to adopt the same fold as the BoNT H_CC_ domain (Fig. 1*B,* Fig. S2). A ricin-type beta-trefoil fold was predicted for the C-terminal region of the putative *C. piperi* toxins by three separate methods (HHpred, Pfam, and Phyre with *E*-values < 0.001). Interestingly, a ricin-type beta-trefoil domain from the *C. botulinum* hemagglutinin, HA33, was identified as the top template by PHYRE (Fig. S2), indicating yet another link between the *C. piperi* toxins and BoNT clusters.

### Genome resequencing of Chryseobacterium piperi confirms presence of bont-like gene clusters

The putative *C. piperi* toxins are located on three separate contigs (2, 44, and 59) in the original draft *C. piperi* genome (NCBI accession JPRJ01, 89 total contigs). To verify a *C. piperi* origin for these contigs, rule out the possibility of sequencing artifacts, and enable further genome-wide analysis, *C. piperi* was acquired from ATCC and sequenced on Illumina MiSeq and Pacific Biosciences RS-II sequencing platforms. A closed genome 4.5Mbp in length, 35.3% GC content, and 250X coverage was produced and analysed (Fig. 3). The assembly revealed one toxin gene cluster (GC1) located at 1399-1432 kbp consisting of two *bont*-like genes, and a second toxin gene cluster (GC2) located at 3287-3312 kbp consisting of one *bont*-like gene (Fig. 3).

**Figure 3.**
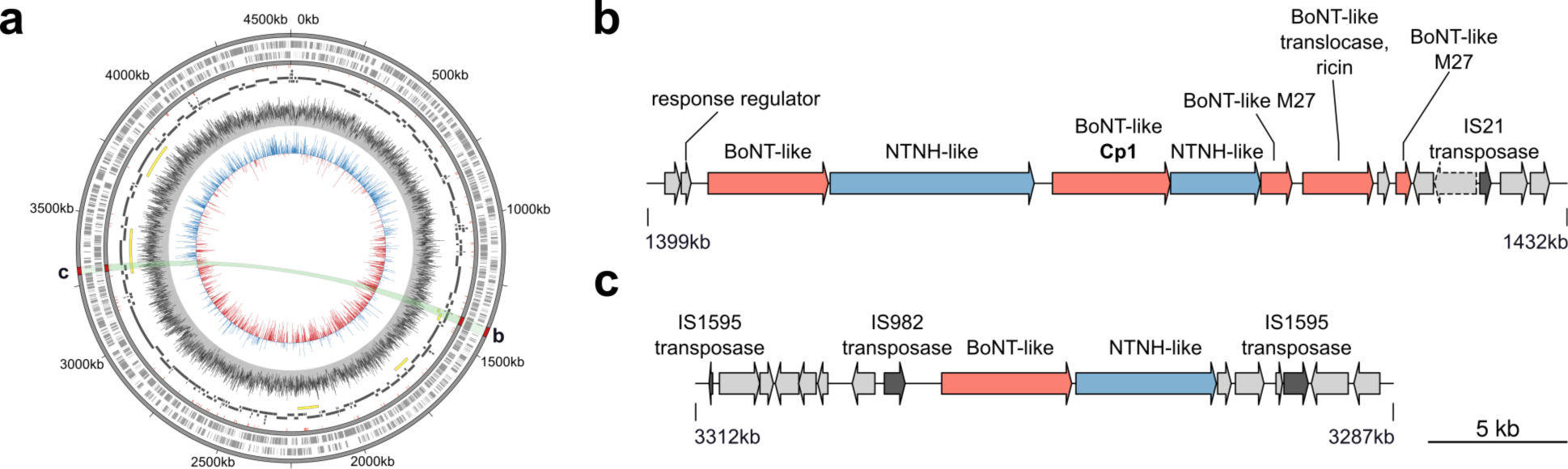
BoNT-like toxins reside within two toxin-rich gene clusters in the genome of *Chryseobacterium piperi*. *A*) Re-sequencing with a combination of Illumina and PacBio resulted in a closed genome with a single circular chromosome. Yellow bands reflect the local alignment of the toxin containing contigs (shown in *B* and *C*) from the initial JPRJ01 assembly against an intermediate assembly (black), and the closed genome (CP023049). Annotated insertion sequences (red ticks), SRX3229522 read mapping to CP023049 (gray histogram), and GC skew (blue/red histogram) are also indicated. The closed genome contains two separate toxin gene clusters (shown in *B* and *C*). Transposase genes are found within and nearby both clusters, which likely contributed to the mobility and duplication of these elements. Full gene annotations for the clusters are available in the Supplementary Information.

Similar to BoNT and BoNT/Wo gene clusters, *C. piperi* BoNT-like toxin genes are flanked by a neighboring gene (designated “NTNH-analog”) with detectable partial homology but lacking the conserved HExxH active site motif. Additionally, the *C. piperi* “NTNH-analog” genes uniquely feature a IBC1 (“Isoprenoid_Biosyn_C1 super family”) domain at the N-terminus similar to class I terpene-synthases whose role is unclear. No additional neurotoxin-associated proteins from the *ha/p47/orfx* families are present in these clusters.

Several genomic features surrounding the *C. piperi* toxin gene clusters indicate an origin via mobile element insertion. First, homologous regions to GC1 and GC2 were not detected in any other available *Chryseobacterium* genomes, suggesting non-*Chryseobacterium* origins. Second, numerous transposases are present including two IS110 family transposases, a IS200/IS605 transposase 18-40 kbp upstream (CJF12_06430, CJF12_06460, CJF12_06500), and an IS1182 family transposase (CJF12_05985) 30 kbp downstream of GC1. IS110 transposases have been previously shown to flank other *bont* gene clusters (44). A disrupted IS982 family transposase pseudogene (CJF12_14555) is located immediately upstream of CJF12_14550 and flanking GC2 are complete and partial IS1595 family, ISChpi, insertion sequences (CJF12_14525 and CJF12_14620). Third, genes neighboring *Chryseobacterium* toxin gene clusters were found to possess homology to genes in *M. chelonae* (e.g., closest homolog of CJF12_14560 was *M. chelonae* WP_064393402.1, 71% amino acid identity), consistent with the detected similarity between the *C. piperi* toxins and *M. chelonae genes* (Fig. 2*A*).

### Chryseobacterium piperi toxin: a novel BoNT-like toxin that induces necrotic cell death

Given the substantial sequence variation between predicted *C. piperi* toxins, we selected WP_034687872 (Cp1) for experimental characterization based on it having the greatest sequence identity to BoNTs (% id = 17%). Initial protease assays of recombinant Cp1 LC against known BoNT substrates, including VAMPs and SNAP25 SNARE proteins, yielded negative results (results not shown). Although the putative toxins from *Chryseobacterium* did not display activity against canonical BoNT targets, the conservation of the active site residues and similarity to M27 and M91 metalloproteases suggested the possibility of other targets. We elected to test for broad, metallopeptidase-induced toxicity via transfection and subsequent expression of the Cp1 LC cDNA in human embryonic kidney HEK293T cell line. CP1-LCs with active site knockouts at H209A and E210A, were utilized as negative controls.

As shown in Fig. 4*A*, the expression of wild-type CP1-LC resulted in a cell death phenotype in HEK293T cells. These cells stopped proliferating and were visibly shrunken, eventually dying and detaching from culture plates. Cell counts after 48 hours revealed an almost 4-fold reduction in the number of cells (Fig. 4*B*). No cell death phenotype or significant reduction in cell number was observed in the H209A and E210A mutants, confirming that the observed toxicity is metalloprotease-dependant. To further confirm the effect of CP1-LC, we performed cell apoptosis assays by flow cytometry using Hoechst 33342, YO-PRO-1, and propidium iodide (Fig. 4*C*). In this assay, live cells (blue), apoptotic cells (blue and green), and dead cells (bright red) are visualized by fluorescence. The percentage of necrotic death was much higher (an approximately 65% increase, see boxed region) in cells transfected with CP1-LC than in cells transfected with control plasmid, H209A, or E210A mutants. In contrast, the percentages of cells labeled as apoptotic death did not change appreciably. These results suggest that expression of CP1-LC leads to necrotic death of cells, and that the cell death depends on the protease activity of CP1-LC. Although the specific target(s) of CP1-LC remains unknown at this point, CP1-LC is clearly an active toxin capable of disrupting cellular processes required for proliferation and survival of human cells.

**Figure 4.**
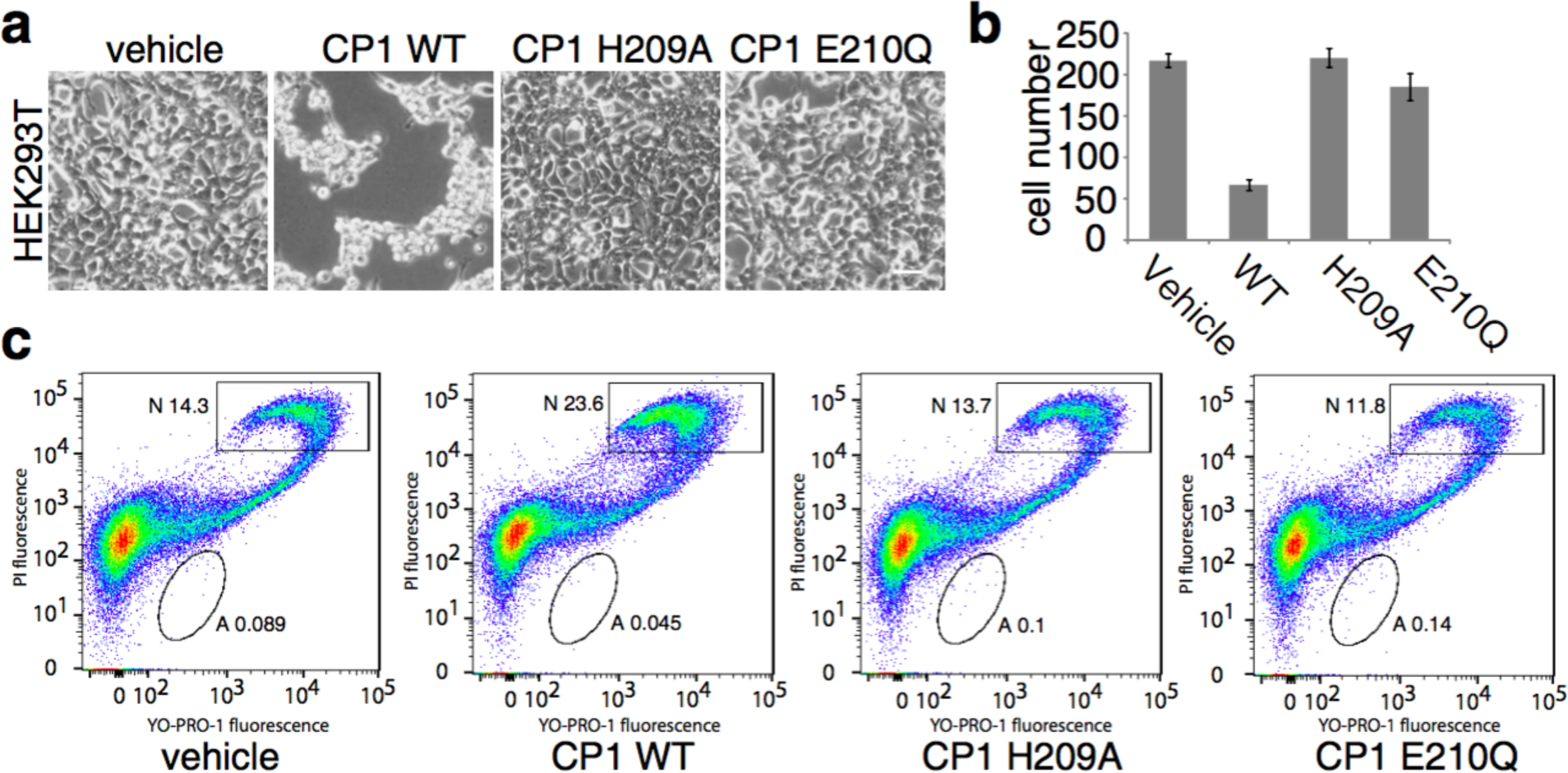
Expression of CP1-LC in HEK293T cell caused necrotic cell death, which depends on its metalloprotease activity. *A*) HEK293T cells were transfected with vehicle vector (pcDNA3.1(+)), CP1-LC WT and mutants (H209A, E210Q) which abolished the metalloprotease activity. Cells were observed and images were taken under inverted microscopy after 48 h. Only cells transfected with CP1-LC WT showed dramatically cell death phenotype which were shrunken and detached from plate. *B*) Cell numbers were counted in defined field of images taken in *A*). *C*) Cell apoptosis assay was carried out with Chromatin Condenstion/Membrane Permeability/Dead cell Apoptosis kit. Transfected cells with vehicle vector, CP1-LC WT and mutant plasmids were analyzed with flow cytometer. Only transfection with CP1-LC WT plasmid enhanced necrotic population of HEK 293T cells which is not observed in inactive form CP1-LC plasmids.

## Discussion

Bacterial toxins such as BoNTs and TeNTs are causative agents of human disease but their evolutionary ancestry likely extends beyond the modern lineages in which they reside. Consistent with this idea, our survey of existing bacterial genomes revealed a diverse family of BoNT-related toxins outside of *Clostridium* including environmental strains of bacteria that are pathogens of non-human hosts (33). These new toxin genes expand the BoNT family and suggest BoNTs are a derived lineage of a broader and potentially ancient toxin family.

Following the recently discovered *Weissella* toxin, *Chryseobacterium* BoNT-like toxins appear to be the next closest evolutionary relatives to BoNTs. The identification of BoNT homologs that cluster outside of the BoNT family (Fig. 2*A*) and possess toxin activity (Fig. 4) raises the strong possibility that BoNTs evolved from precursor proteolytic toxins targeting different substrates. By assessing the domain architectures of BoNT-like toxins in the context of the phylogenetic tree, it is possible to reconstruct a stepwise model of BoNT evolution (Fig. 5). This phylogenetic model suggests that BoNTs evolved from an ancestral toxin containing a M27-like protease domain, which in turn is related to the more distant M91 family of single domain type III effector proteases. The ancestral toxin also likely possessed a translocation domain that may have more closely resembled the T domain of diphtheria toxin. Finally, the ancestral toxin also likely possessed a receptor-binding region encoding a beta-trefoil ricin-type fold. Evolutionary novelties that occurred in the lineage leading to BoNTs appear to include extension of the translocation domain to include the large alpha-helical bundle, the gain of the N-terminal LamG-like binding domain, as well as the acquisition of toxin accessory genes neighbouring the proteolytic toxin gene (including NTNH and HA or P47/ORFX proteins). The increased sequence diversity of BoNT-like toxins, as well as the taxonomic diversity of their hosts, suggests that these represent early-branching lineages in BoNT evolution, and the canonical domain architecture of BoNTs has emerged more recently.

**Figure 5.**
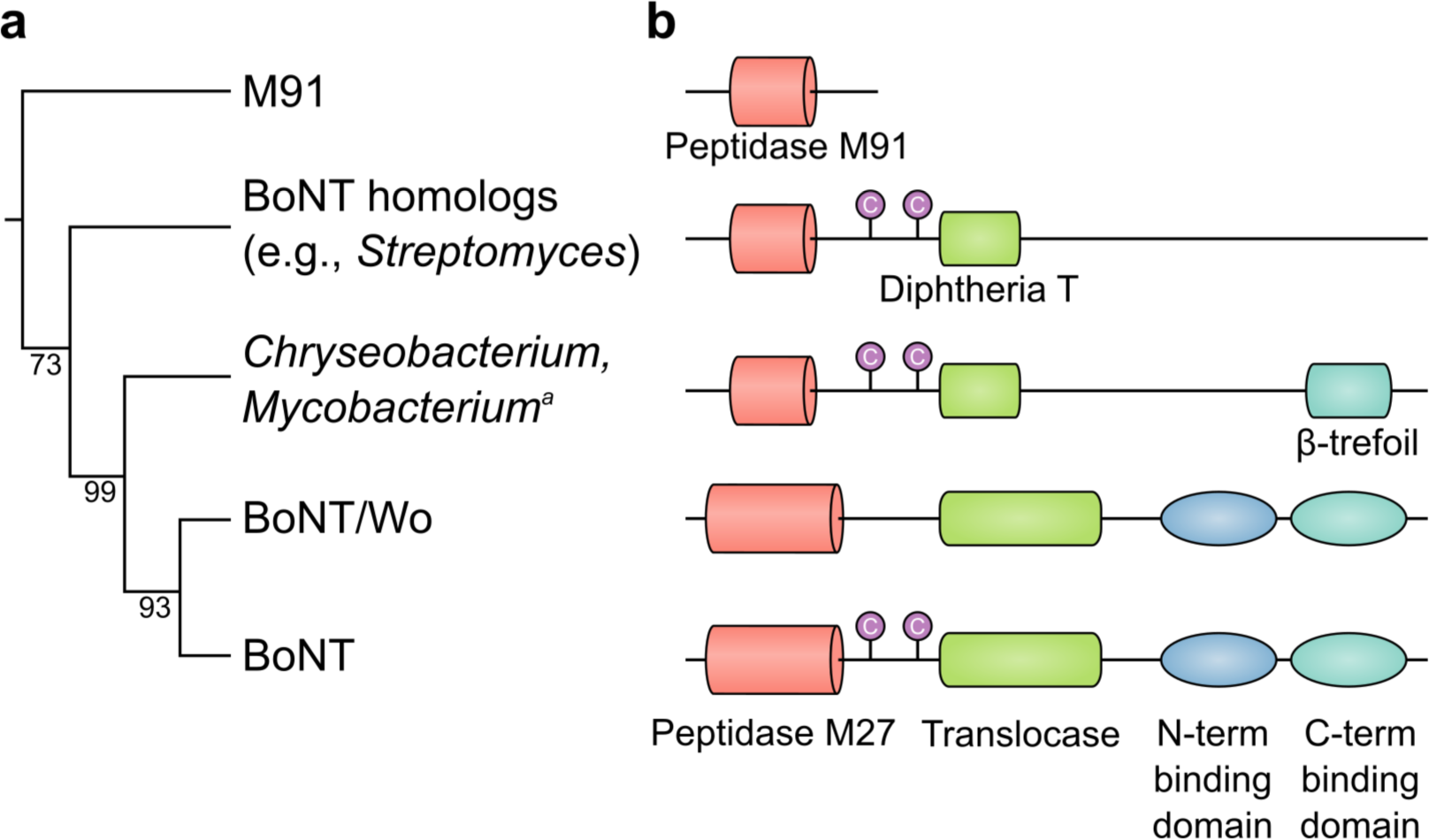
A model of BoNT evolution from ancestral toxins. *A*) Simplified cladogram based on an alignment of the BoNT peptidase domain (the most conserved region in BoNTs). *B*) Domain architecture changes over BoNT family tree. In the near-BoNT clades, all BoNT domains are detectable. Other more ancestral BoNT homologs lack identifiable BoNT domains, highlighting the divergent nature of the BoNT family.

The future identification of the substrates targeted by *Chryseobacterium* toxin, *Weissella* toxin and others, combined with determination of their structure, will illuminate the specific evolutionary changes in the BoNT LC responsible for its gain of activity against neuronal SNAREs. We anticipate that this work will ultimately uncover a detailed model for the origin of the most deadly toxin known.

## Methods

### Detection, comparison and analysis of bont-like genes

Sequences were retrieved using PSI-BLAST with default parameters (BLOSUM62 scoring matrix; expect threshold 10; gap open 11; extension 1) from the nr database on (March 26, 2017) (45). Initial homologs were discovered by searching with BoNT/A1 (NCBI accession number ABS38337.1) with up to two rounds of PSI-BLAST. Then, in order to retrieve all possible sequences from each sequence family, different queries were used to search for specific BoNT homolog subfamilies (*Chryseobacterium*: WP_034681281.1, *Actinobacteria*: *Streptomyces* sp. NBRC 110027 GAO13068.1, fungal: *Metarhizium anisopliae* KFG81441.1) and reiterating to convergence. BoNT homologs identified this way were added to a set of known BoNT and NTNH proteins represnting all known serotypes, including the recently discovered BoNT/F5A (KGO15617.1) and BoNT/X (KGO12225.1). Sets of T3SS effectors and diphtheria toxins were also retrieved via PSI-BLAST, with diphtheria toxin (PDB accession number 4AEO.1) and *E. coli* NleD (WP_069191536.1) as the original queries. These sets of T3SS effectors and diphtheria toxins were pruned to remove identical sequences using Jalview (46). To obtain the *Austwickia* pseudogene corresponding to the diphtheria C domain, we translated the region immediately upstream of the protein containing diphtheria T and R domains (the region 39931-43386 on contig NZ_BAGZ01000024.1 of *A. chelonae* NBRC 105200).

All-by-all sequence pairwise alignments were generated with needle (of the EMBOSS package, v6.6.0.0 (47)) with default parameters (gap open = 10, gap extend = 0.5, EBLOSUM62 scoring matrix). In Figure 1, percent similarity was used over percent identity in order to allow divergent homologs to cluster more accurately. Principal coordinate analysis was performed in R on a distance matrix of pairwise similarity values using the default dist() and cmdscale() functions.

Domains were annotated with hmmscan (v3.1b2, available from http://hmmer.org/) against the Pfam database v31.0 (48) with an *E*-value cutoff of 1e-6. Annotations were subsequently confirmed by comparison to the Conserved Domain Database with relaxed cutoffs v3.16 (49), and alignment to BoNTs. Full domain annotations are available upon request. For Figure 1, the BoNT homologs with the most BoNT-like annotations were depicted to facilitate comparison between categories.

### Comparison of proteases from BoNTs, BoNT-like proteins and T3SS effector peptidases

All BoNT homologs possessing a putative peptidase domain (i.e., possessing an HExxH motif) were aligned with BoNT and M91 peptidases using Clustal-Omega with defaults v1.2.1 (50), manipulated and colored in Jalview (46). Only regions corresponding to the peptidase domain boundaries were used, the positions of which were estimated based on alignment with domain boundaries of BoNT/A1 (PDB structure 3BTA). The same alignment procedure was used to identify the putative translocation region of BoNT homologs (Fig. S6). After identifying putative domains in BoNT homologs, the segments were combined and realigned.

A maximum likelihood phylogeny for BoNT, BoNT homolog and T3SS peptidases was generated using RAxML (v8.2.4, (51)) (see simplified cladogram in Fig. 5, for the full tree see Fig. S4). Bootstrap support was calculated using 100 rapid bootstraps.

### Structural modelling of BoNT-like proteins and T3SS effector peptidases

Structural templates were identified for Cp1 (*C. piperi*, accession WP_034687872.1) using the LOMETS meta-server (52) on July 18, 2017. Templates (PDB IDs: 3BTA:A, 1XTG:A, 5BQN:A) were selected based on highest significant threading alignments (normalized Z-scores: 5.12-1.21, identity: 17%-21%). Structural modelling and refinement was done through ITASSER (53), and the model with the lowest C-score was selected. For *E. coli* NleD, structural templates were identified through GeneSilico Metaservers (PDB IDs: 1Z7H:A, 1EPW:A, 3BWI:A, 3DEB:A, 3BON:A, 2QN0:A, 2A97:A, 3DDA:A, 1XTG:A, 1ZB7:A, 1F0L:A, 3FFZ:A, 1YVG:A, 2FPQ:A, 2G7K, 5BQN:A, 2NYY:A, 1T3C:A, 3V0A:A, 3FIE:A, 1RM8:A, 1E1H:A, 3VUO:A, 2A8A:A, 3D3X:A, 3DSE:A) were selected from COMA (score ≤ 5.4e-07, identity: 19%), HHblits (score: 100, identity: 13%-20%), and HHsearch (score: 96.3, identity: 13%-19%) on July 15, 2017. Structural modelling was carried out through PRIMO's pipeline (54). Identified template sequences were aligned to M91 with T-Coffee Expresso (55), which uses 3D-Coffee to incorporate structural information during alignment. A total of 20 homology models were created with slow refinement based on the resulting alignment using MODELLER (56). The model with the lowest DOPE Z-Score was selected. Structural quality was assessed with Ramachandran plot analysis using PROCHECK (57). Models were visualized using Chimera (58).

### Re-sequencing of the Chryseobacterium piperi genome

Methods, materials, and platforms used in the sequencing and assembly of *C. piperi* are described in Wentz et al. (59). The closed genome is accessible at the DDBJ/ENA/Genbank under the accession number CP023049. MiSeq and RS-II reads utilized in assembly are available at NCBI SRA under accessions SRX3229522, SRX3231351, SRX3231352. Figure 3 was generated using the program Circos (60).

### HEK293T cell transfection and cell number counting

HEK 293T cells were dispensed on 24-well plate at the density of 0.2× 10^6^ cells/well. After 24 h, cells were transfected with 0.5 *μ*g vehicle vector (pcDNA3.1(+)), CP1-LC WT(1-398 aa, NCBI accession number WP_034687872.1), CP1-LC H209A and CP1-LC E210Q plasmids with PolyJet reagent. Pictures were taken 48 h later after transfection. Cells numbers were counted in defined field (6× 6 inch) from three different pictures.

### HEK293T cell death assay

HEK293T cells were dispensed on 60 mm dish at the density of 2.5× 10^6^ cells/dish. Cells were transfected with 2.5 *μ*g vehicle vector (pcDNA3.1(+)), CP1-LC WT, CP1-LC H209A and CP1-LC E210Q plasmids by using 5 *μ*L PolyJet. Cells were harvested 24 h after transfection and washed with cold phosphate-buffered saline (PBS). Cell density was adjusted to 1× 10^6^ cells/mL. A mixture of 1*μ*L Hoechst 33342, YO-PRO-1 and propidium iodide stock solution (Invitrogen) were added into a 1 mL cell suspension. Cells were incubated on ice for 30 minutes. Stained cells were analyzed by flow cytometry (BD/Cytek FACSCalibur DxP 11). UV excitation was used for detection of 460 nm emission of Hoechst 33342 dye, 488 nm excitation was used for detection of the 530 nm emission of YO-PRO-1 dye, and 575 nm emission of propidium iodide. Cell population separated into three groups: live cells showed only a low level of blue fluorescence, apoptotic cells showed bright green and blue fluorescence and dead cells showed bright red fluorescence.

## Acknowledgments

We thank Nagarajan Thirunavukkarasu, Sara Lomonaco, and Tim Muruvanda for their technical advice and useful discussions.

## Author Contributions

A.C.D., S.K.S. and M.D. conceived and coordinated the project. T.G.W. and S.K.S. performed genome sequencing. M.J.M, T.G.W., and A.C.D. performed bioinformatic analyses. S.Z. performed all biochemical analyses. All authors contributed to writing of the manuscript.

